# Structures of virus-like capsids formed by the Drosophila neuronal Arc proteins

**DOI:** 10.1101/697193

**Authors:** Simon Erlendsson, Dustin R. Morado, Jason D. Shepherd, John A. G. Briggs

## Abstract

The neuronal protein Arc is a critical mediator of synaptic plasticity. Arc originated in tetrapods and flies through domestication of retrotransposon *Gag* genes. Recent studies have suggested that Arc mediates intercellular mRNA transfer and like Gag, can form capsid-like structures. Here we report that Drosophila proteins dArc1 and dArc2 assemble virus-like capsids. We determine the capsid structures to 2.8 Å and 3.7 Å resolution, respectively, finding similarity to capsids of retroviruses and retrotransposons. Differences between dArc1 and dArc2 capsids, including the presence of a structured zinc-finger pair in dArc1, are consistent with differential RNA-binding specificity. Our data support a model in which ancestral capsid-forming and RNA-binding properties of Arc remain under positive selection pressure and have been repurposed to function in neuronal signalling.

## Introduction

The immediate early gene *Arc* is a master regulator of synaptic plasticity (Shepherd and Bear, 2011), essential for consolidation of memory (Plath et al., 2006) and experience-dependent long-lasting changes in the mammalian brain (McCurry et al., 2010). *Arc* transcription is rapidly induced by experience through neuronal activity and *Arc* mRNA is trafficked out to dendrites (Link et al., 1995; Lyford et al., 1995; Steward and Worley, 2001) where it is locally translated (Waung et al., 2008). Arc protein regulates synaptic function via trafficking of AMPA-type glutamate receptors (Chowdhury et al., 2006; Shepherd et al., 2006) and interacts with proteins at the synapse including PSD95 (Fernandez et al., 2017; Okuno et al., 2012), the transmembrane AMPAR regulatory protein *γ*2 (TARP*γ*2) (Jackson and Nicoll, 2011), Ca^2+/^calmodulin dependent kinase (CaMKII) (Okuno et al., 2012) and several proteins involved in clathrin-dependent endocytosis (Chowdhury et al., 2006; DaSilva et al., 2016). Thus, elucidating the structure and function of Arc will shed light on fundamental mechanisms of cognition.

The *Arc* gene is present throughout the tetrapods, but is not found in fish or lower vertebrates. An Arc homologue, however, is found in the fly lineage, where it is involved in synaptic plasticity at the neuro-muscular junction (Ashley et al., 2018; Pastuzyn et al., 2018). *Drosophila melanogaster* contains two copies of this Arc homologue (dArc1 and dArc2). Based on phylogenomic analyses, tetrapod Arc and fly Arc originate from two independent domestication events from members of the Ty3/Gypsy family of Long Terminal Repeat (LTR) retrotranspons (Abrusán et al., 2013; Campillos et al., 2006; Pastuzyn et al., 2018). LTR-retrotransposons are self-replicating genetic elements that integrate into the host’s genomic DNA. Active and inactive retrotransposons make up large fractions of the genomes of multi-cellular organisms (Lander et al., 2001; Venter et al., 2001). The group specific antigen (Gag) domain in active Ty3/Gypsy retrotransposons encodes the CA protein that forms a virus-like protein capsid with extensive structural homology to that of enveloped retroviruses such as Human Immunodeficiency Virus (HIV) (Dodonova et al., 2019; Nielsen et al., 2019; Zhang et al., 2015).

Remarkably, recent studies show that both tetrapod Arc and dArc1 may have preserved the ability to form capsid-like structures, and that secreted extracellular vesicles mediate transfer of Arc protein and mRNA between cells (Ashley et al., 2018; Pastuzyn et al., 2018). These observations suggest that Arc capsids may package and transfer their own mRNA between cells, providing a new mechanism for intercellular communication. Here, we report the structures and molecular architecture of the dArc1 and dArc2 capsids. dArc1 and dArc2 form icosahedral capsids, with structural similarity to those of Ty3 and mature HIV-1, showing that endogenous proteins that function in neuronal-signaling form virus-like capsids.

We recombinantly expressed dArc1 and dArc2 in *Escherichia coli* and purified both proteins using heparin affinity and an N-terminal Glutathione-S-Transferase (GST)-tag. Removal of the GST-tag led to assembly of homogenous, spherical particles with outer diameters of ∼37 nm and a dense capsid layer of ∼3 nm in thickness (Fig. 1A,D). Using single particle cryo-EM we determined the structures of dArc1 and dArc2 capsids at 3.5 Å and 3.9 Å resolution, respectively (Fig. 1 and S1, Table S1), and by performing symmetry expansion and local refinement, improved the resolution to 2.8 Å and 3.7 Å, respectively (Fig. S2, S3, Table S2). dArc1 and dArc2 assemble icosahedral capsids with triangulation number T=4 and are composed of 12 five-fold symmetric pentameric capsomeres and 30 two-fold symmetric hexameric capsomeres (Fig. 1). Short spikes (5-8 nm long) protrude from the center of each capsomere.

**Figure 1:**
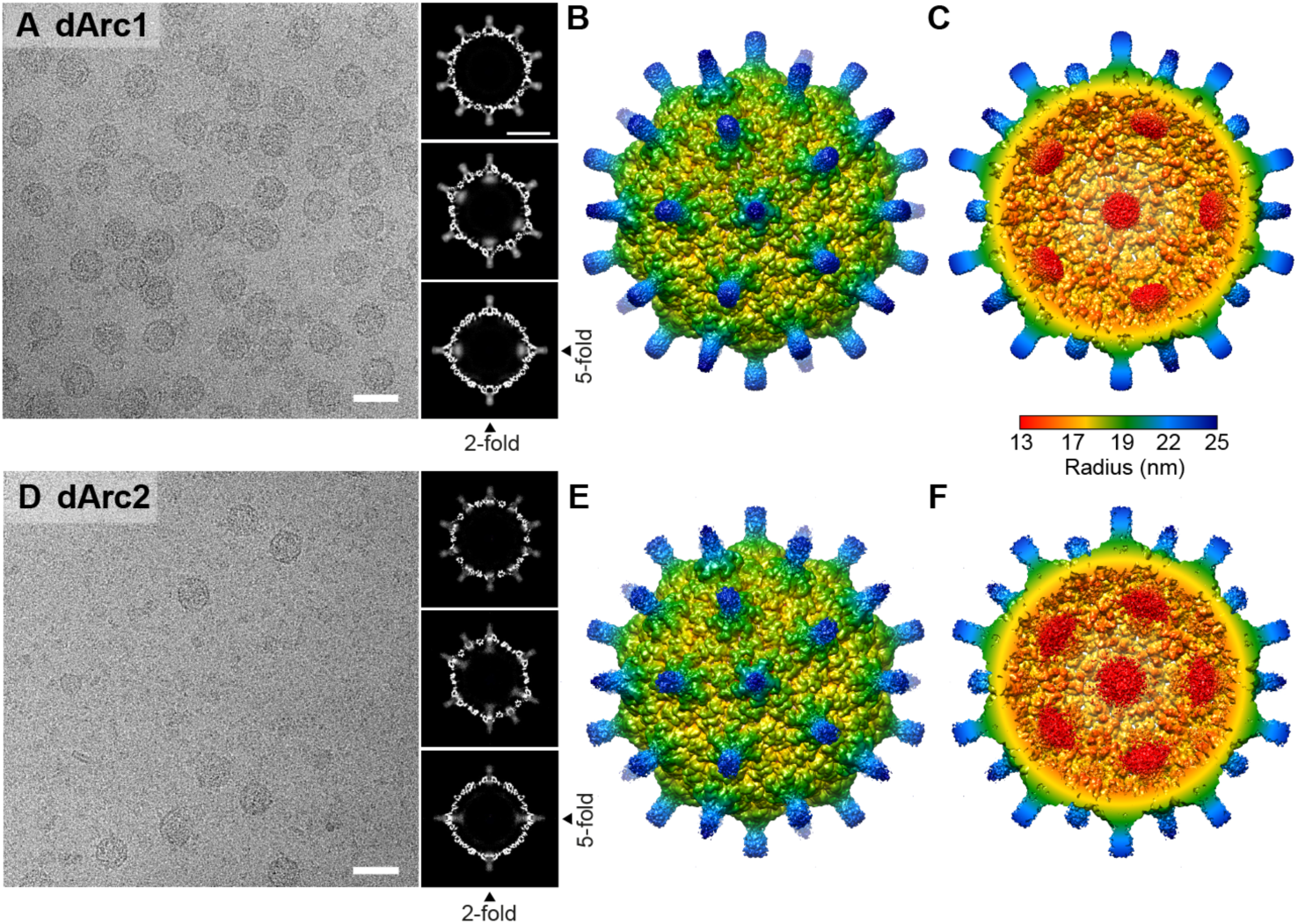
The cryo-EM structures of dArc1 and dArc2. **A)** Representative dArc1 micrograph and central sections through three-dimensional reconstructions perpendicular to the five-, three-, and two-fold axes. Scale bars are 50 nm for the micrographs and 20 nm for the slices. **B**) Surface representation of dArc1, viewed down the icosahedral five-fold axis and colored by radius. **C**) As in B, with the front half of the capsid removed to real internal features. **D-F)** As in A-C, but for dArc2.

We built atomic models for the capsids into the structures obtained for assembled dArc1 and dArc2 (Fig. 2 and S4A, Table S3). The capsid shells are formed by 240 copies of the predicted CA domain (dArc1 residues 42-205, dArc2 residues 29-192) with four independent copies of CA in the asymmetric unit (Fig. 2A-B). The dArc1 and dArc2 CA structures are highly similar, with RMSDs <0.8 Å for the individual CA molecules in the asymmetric unit, reflecting their similar sequences (Fig. S6). CA consists of two connected globular domains, CA_NTD_ and CA_CTD_, connected by a flexible central linker (Fig. 2B). The CA_NTD_ (dArc1 residues K42-S120) constitutes the central part of each capsomere (Fig. 2C-D), which consists of four *α*-helices, *α*1-*α*4, and an extended chain (dArc1 residues 42-55). The CA_CTD_ (dArc1 residues A125-H203) forms the “valleys” between protruding capsomeres and consists of five *α*-helices, *α*5-*α*9 (Fig. 2B-D).

**Figure 2:**
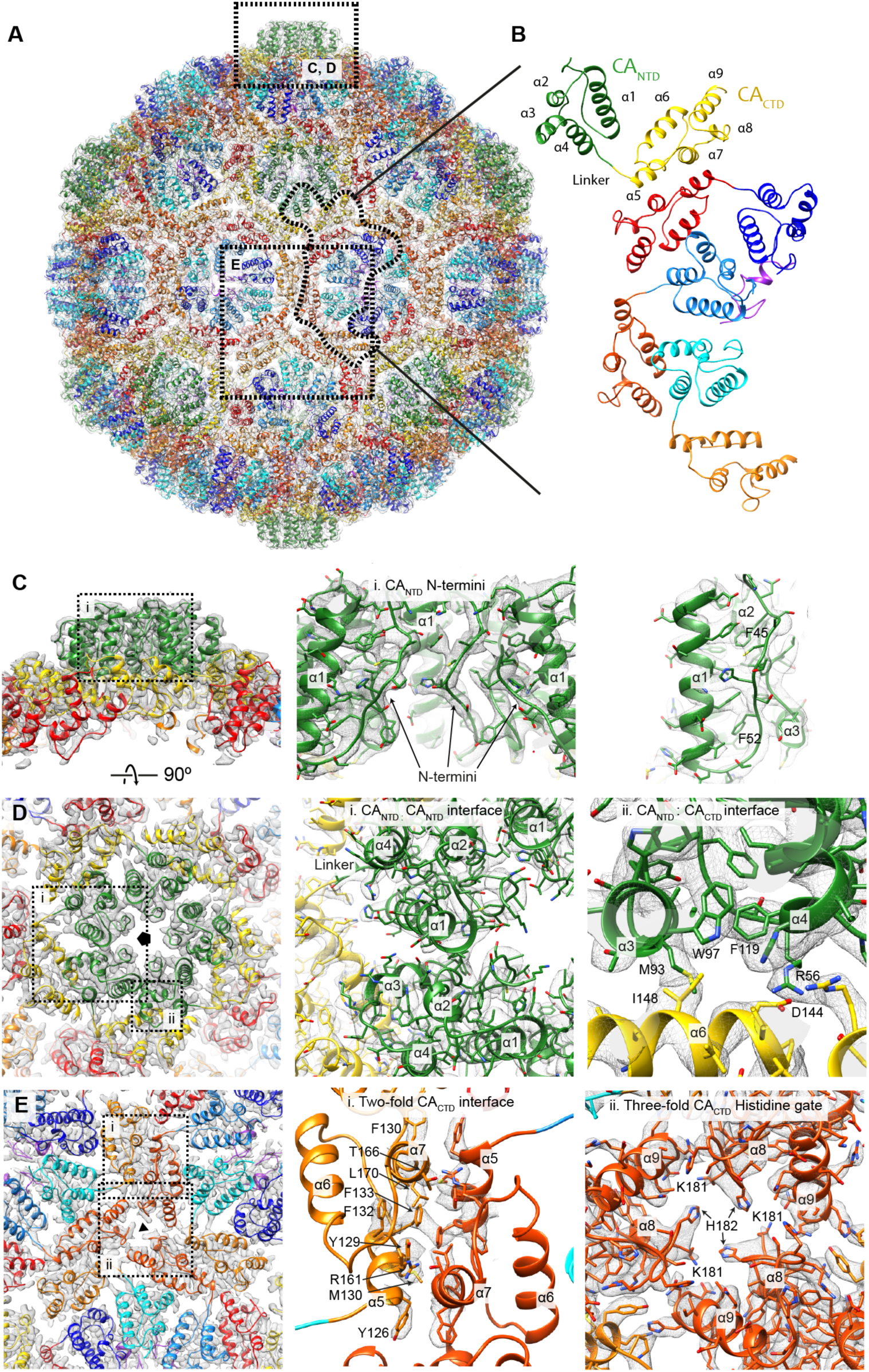
Atomic model of the dArc1 capsid. **A)** The 12 five-fold capsomeres are colored green (CA_NTD_) and yellow (CA_CTD_). The 30 two-fold capsomeres are colored cyan to blue (CA_NTD_) and orange to red (CA_CTD_). The zinc fingers are colored purple. EM density is transparent grey. **B)** The asymmetric unit containing four CA molecules and one zinc finger. 60 asymmetric units including 240 individual CA molecule and 60 zinc fingers make up the T=4 capsid. **C)** Close-up of the five-fold capsomere (outlined in A) i) Cut-away showing three out of five N termini in the centre of the capsomeres. The N-termini extend into and form the capsid spikes. The N termini are stabilized by docking into an extended hydrophobic groove adjacent to *α*1. **D)** External view of the five-fold capsomere i) The CA_NTD_:CA_NTD_ interaction between *α*1, and *α*2 and *α*3 of the neighbouring CA molecule in the capsomere. ii) The CA_NTD_:CA_CTD_ interface which involves *α*6 in the CA_CTD_ and *α*3 and *α*4 in the neighbouring CA_NTD_. This interface relies on both hydrophobic and electrostatic interactions. **E)** External view of the two and three-fold CA_CTD_ interfaces. i) The two-fold CA_CTD_ interface connects two adjacent capsomeres and is dominated by hydrophobic π-π stacking interactions. The interface involves residues from *α*5 and *α*. EM density is shown only for the contact area of the interface. All CA_CTD_ interfaces are highly similar (Fig. S7). ii) The three-fold CA_CTD_ axis. At this position, basic residues from *α*8 surround the largest gap in the capsid. The corresponding views for dArc2 are shown in Fig. S4.

The structures of the dArc1 CA_NTD_ and CA_CTD_ domains are similar to those of monomeric rat Arc (rArc) in solution (Nielsen et al., 2019) (RMSDs of 1.3 and 1.2 Å, respectively; Fig. S5A-B), though the linker connecting CA_NTD_ and CA_CTD_ is more rigid in monomeric rArc. The protein-protein interfaces in the dArc capsid do not correspond to those predicted based on rArc crystal contacts (Zhang et al., 2019). The dArc extended N-terminal chain (dArc1 residues 42-55, dArc2 residues 29-32) binds into, and occludes, a hydrophobic groove between *α*1 and the linker between *α*2 and *α*3 (Fig. 2C), which is the binding site for TARP*γ*2, CaMKII and the NMDA receptor peptides in monomeric rArc (Nielsen et al., 2019) (Zhang et al., 2015) (Fig. S5C). These observations are consistent with a requirement for domain reorganization or uncovering of one or more interaction surfaces to initiate rArc CA oligomerization (Nielsen et al., 2019).

The CA molecules within the dArc capsomere are held together by two distinct interfaces: (i) the CA_NTD_:CA_NTD_ interface is formed by acidic residues from *α*1 and basic residues from the neighboring *α*2 and *α*3 (Fig. 2D); (ii) the CA_NTD_:CA_CTD_ interface involves docking of I148 from CA_CTD_ *α*6 into a hydrophobic pocket created between *α*3 and *α*4 from the neighboring CA_NTD_, and formation of a potential salt bridge between D144 in *α*6 and R56 in *α*1 (Fig. 2D). The capsomeres are then joined together by a dimeric CA_CTD_:CA_CTD_ interface, dominated by hydrophobic and *π*-stacking interactions between adjacent *α*5 and *α*7 (Fig. 2E.i). CA_CTD_ molecules also interact around the three-fold symmetry axis between the hexameric capsomeres and around the pseudo three-fold symmetric axis between two hexameric and one pentameric capsomere(s). In both dArc1 and dArc2, the three-fold and pseudo-three-fold CA_CTD_ axes provides the biggest accessible openings in the capsid surface. At these positions in dArc1, the sidechains of K181 and H182 extend towards the center of the opening and are positioned 6 or 10 Å apart (Fig. 2E.ii and S7H). In dArc2, it is the sidechains of E168 and S169 that extend towards the center of the opening (Fig. S4E.ii). We speculate that changes in the protonation state of H182 in dArc1 could modulate capsid stability or the transport of ions in or out of the capsid.

The four independent copies of CA in the asymmetric unit differ only in the relative orientation of CA_NTD_ and CA_CTD_ (Fig. S7D). The interfaces formed by independent copies of the CA_CTD_ are similar to one another (Fig. S7G) and to those present in the Ty3 and mature HIV-1 capsids (Fig. S8, S9). In contrast, the CA_NTD_-CA_NTD_ interface differs among the four independent copies of CA (Fig. S7E), and between dArc and Ty3/Gypsy (Fig. S9F-G). This reflects the divergence of CA_NTD_ structure and arrangement among retrotransposons and retroviruses, perhaps due to the role of the CA_NTD_ in interacting with divergent host factors (Schur et al., 2015; Zhang et al., 2015).

dArc1 contains 41 amino acids upstream of CA_NTD_ (Fig. S6), while dArc2 has a 28 amino acid upstream region similar to that of dArc1 except for the absence of a poly-A stretch (Fig. S6). These amino acids form the exposed spikes. The spikes are highly flexible, and their structures are therefore not resolved (Fig. 1, Fig. S3). There are six copies of the N-terminal region above the two-fold axis, but only five copies at the five-fold axis. The N termini have predicted propensity to form amphipathic *α*-helices (Fig. S10), the hydrophobic face of which is likely to mediate oligomerization or binding to cellular proteins or membranes. Tetrapod Arc has an extended ∼200 amino acid N-terminal domain that may bind RNA (Eriksen et al., 2019). In dArc, the N terminus occludes openings in the CA shell at the five-fold and two-fold axes. Thus, access to the capsid interior through these openings could by regulated by N-terminal interactions.

Interestingly, we observe a disordered/diffuse density inside the capsid beneath the five-fold axis capsomeres for both dArc1 and dArc2 capsids (Fig. 1), although the dArc2 protein sequence terminates immediately at the end of *α*9 of CA_CTD_ (Fig. 3, Fig. S6). This suggests that some copies of the N-terminal region constitute the spike, while others protrude inwards to form this density.

**Figure 3:**
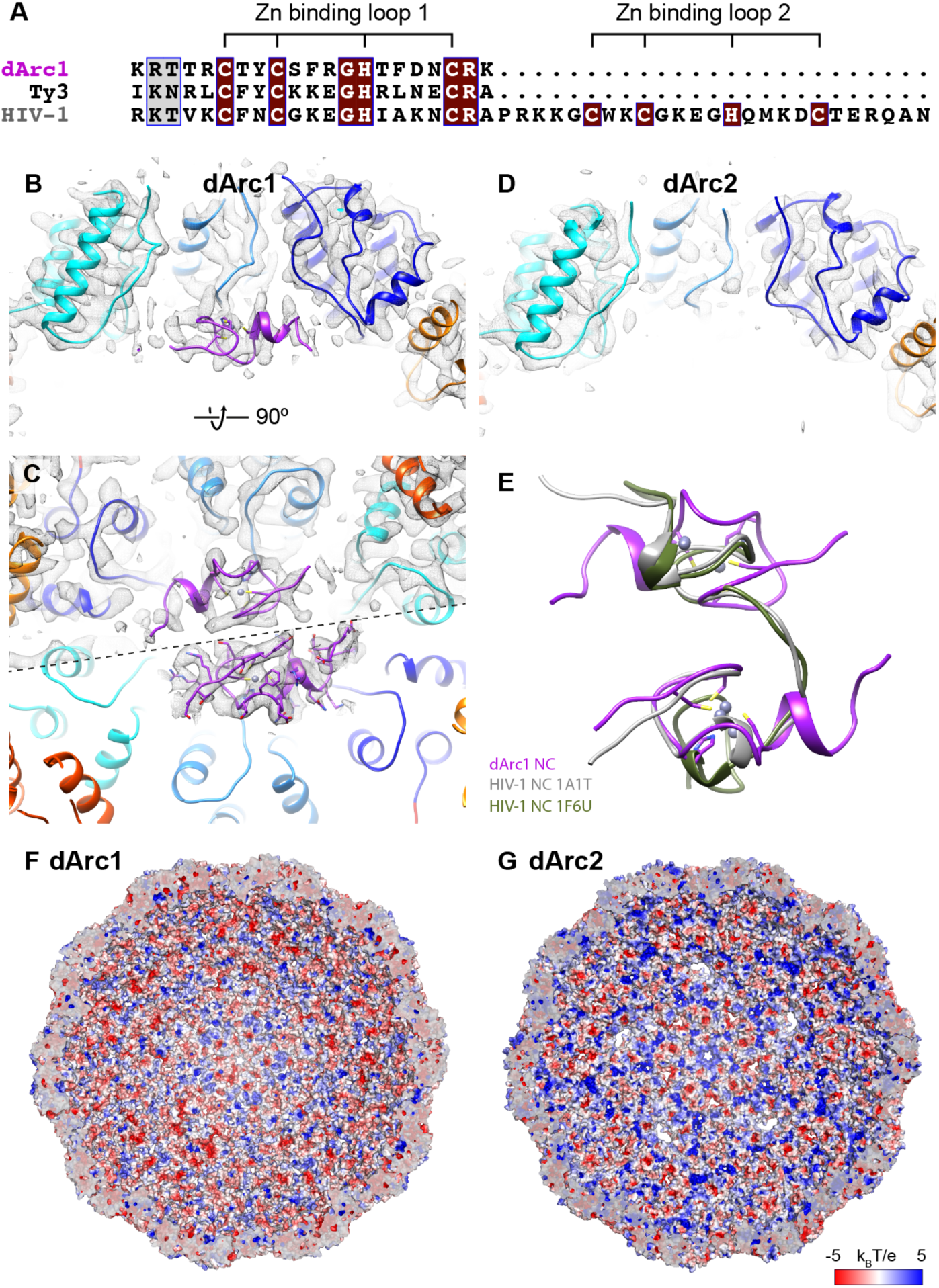
Residues within the dArc1 NC domain form a zinc finger bound to CA. **A**) sequence alignment between the zinc finger regions of dArc1, Ty3 and HIV-1. **B)** Cross-section of the two-fold capsomere showing the fit and position of the zinc finger (purple). **C)** Viewed from the inside of the capsid, two zinc-fingers are seen bound beneath the two-fold capsomeres. The bottom half of the panel shows EM density for the zinc finger at a lower isosurface threshold. **D**) dArc2 lacks the NC domain, and EM density for the zinc finger is absent. **E)** overlay of the two copies of the dArc1 zinc fingers at the two-fold capsomere (purple), and the HIV-1 double zinc finger motif in the SL2-bound (green) and SL3-bound (grey) configurations. **F-G**) Cross-sections of the entire dArc1 and dArc2 capsids surfaces colored by electrostatic potential from -5 (red) to +5 k_b_T/e (blue).

dArc1 has 48 additional residues downstream of CA_CTD_ (dArc1 NC), that extend into the capsid interior and are homologous to the nucleocapsid (NC) domains of retroviruses/retrotransposons (Fig. 3A), which mediate specific nucleic acid interactions. Below the two-fold axis, two copies of residues 224-252 in dArc1 NC are well resolved, forming single-knuckle zinc fingers, arranged in an anti-parallel manner (Fig. 3C-D). The zinc fingers are absent in dArc2. Similar to the two zinc fingers in the HIV-1 NC, the zinc molecule is tetrahedrally coordinated by three cysteines and one histidine residue (C227, C230, H235 and C240). The zinc fingers are connected to one another by salt bridges between R226 and Q248. This orientation of the zinc fingers guides the majority of basic residues (R233, 241, 247 and K245) into electronegative patches created in the interface between the *α*1 and the adjacent *α*3 in CA_NTD_. Strikingly, the spatial separation of the two zinc fingers from adjacent copies of dArc1 NC is similar to the separation of the two zinc fingers within a single copy of HIV-1 NC when bound to SL2 or SL3 in the HIV-1 genomic packaging signal (Amarasinghe et al., 2000; De Guzman et al., 1998)(Fig. 3E). This arrangement of zinc fingers facilitates binding of HIV-1 NC to three exposed bases in the tips of these stem loops.

The hydrophobic cores of dArc1 and dArc2 CA domains are highly conserved in sequence and structure. Protein-protein interfaces in the capsid are also highly-conserved: out of the 36 more radical amino acid substitutions, 31 are exposed, and only four are at protein-protein interfaces, all of which are at the CA_NTD_:CA_NTD_ interface. Notably, 10 of the amino acid differences between dArc1 and dArc2 involve additional basic residues in dArc2 that are dispersed in linear sequence but are located on the surface of the capsid. Consequently, the interior surface of the dArc2 capsid is more electropositive than dArc1 (Fig. 3F,G, Fig. S11), while dArc1 has additional basic residues in the disordered regions of NC. The basic character of capsid interiors of both dArc1 and dArc2 would facilitate non-specific electrostatic packaging of RNA, while the zinc finger unique to dArc1 may facilitate specific recognition of the 3’ UTR of dArc1 mRNA that is absent in dArc2. Consistent with this observation, dArc1 mRNA is highly enriched and much more abundant in extracellular vesicles than dArc2 mRNA, which lacks a long 3’UTR (Ashley et al., 2018).

The Ty3 retrotransposon packages two copies of its 5.2 kb genome into a T=9 capsid with an internal volume of ∼5 × 10^4^ nm^3^. The T=4 dArc1 capsid is much smaller, with a volume of ∼1.7 × 10^4^ nm^3^. It would enclose RNA at a similar density to Ty3 if it packaged two copies of the ∼2.3 kb dArc1 mRNA (including the 3’ UTR). We suggest that a shrinkage of the capsid from T=9 to T=4 for dArc occurred post-domestication, reflecting shortening of the packaged RNA from a full length LTR retrotransposon mRNA (containing genes coding for both the Gag protein and the RNA polymerase) to the shorter dArc1 mRNA.

Our observations provide a structural basis for future experiments to modulate the function of Arc capsids in synaptic plasticity, or to adapt them to deliver alternative cargos. Taken together, our observations demonstrate that dArc1 and dArc2 have conserved capsid assembly and RNA-packaging features from the ancestral retrotransposon and suggest that these features remain under selection pressure in the host genome. This supports the model that tetrapods and flies have repurposed a retrotransposon gene to package and transfer RNA between cells within virus-like capsids.

## Author contributions

S.E., J.D.S., and J.A.G.B. designed the project. S.E. prepared all samples and performed cryo-EM sample preparation/screening, reconstruction and model building with assistance from D.R.M and J.A.G.B. S.E., J.D.S., and J.A.G.B. prepared the manuscript. D.R.M commented on the final manuscript.

## Acknowledgements

We thank Sjors Scheres, Takanori Nakane and Garib Murshudov (MRC-LMB) for advice on data processing and analysis, and Jake Grimmett and Toby Darling (MRC-LMB) for support with computing infrastructure (MRC-LMB). This study was supported by the MRC-LMB EM Facility. This work was funded by the Novo Nordisk Foundation (S.E: NNF17OC0030788), the National Institute of Mental Health (J.D.S: R01-MH112766) and the Medical Research Council (J.A.G.B. MC_UP_1201/16).

## Materials and Methods

### dArc1 and dArc2 expression and purification

DNA sequences corresponding to dArc1 residues 2-254 and dArc2 residues 2-193 were expressed as Glutathione-s-Transferase (GST) fusion constructs by subcloning into the pGEX 4T1 vector (GE28-9545-49). The plasmids were transform into E. coli BL21(DE3)/pLysS and protein was expressed in autoinduction medium (10 g tryptone, 5g yeast extract, 5 g glycerol, 0.5 g glucose, 2 g α-lactose, 3.3 g (NH_4_)_2_SO_4_, 6.8 g KH_2_PO_4_, 7.1 g Na_2_HPO_4_, 1 mM MgSO_4_, 50 μM ZnCl_2_ per liter - ampicillin and chloramphenicol for selection). Cultures are grown to an OD_600_ of 0.6-0.8 at 37 °C and then shifted to 19 °C for autoinduction overnight (12-16 hrs.). After autoinduction cells were pelleted at 6000 rpm (Rotor JLA 8.100), for 15 min. at 4 °C and resuspended in Lysis buffer (50 mM Tris, 400 mM NaCl, 5 % Glycerol, 2 mM DTT, 50 μM ZnCl_2_, 0.2 mM PMSF (Phenylmethanesulfonyl fluoride), pierce protease inhibitor cocktail (Thermo Scientific A32963), Nuclease Mix (GE80-6501-42) pH 8) and snap frozen in liquid nitrogen. The thawed lysates were sonicated and centrifuged at 16000 rpm (Rotor JA 25.50) for 30 min. The resulting supernatants were cleared by filtration (0.22 μM), and the conductivity of the solution was adjusted to 20 mS/cm before applying it to a Heparin Sepharose column (GE17-0407-01), and eluting using a NaCl gradient. The sample was immediately applied to a GSTrap column (GE17-5282-01) and eluted in TBS (50 mM Tris, 150 mM NaCl, 2 mM DTT, 50 μM ZnCl_2_, pH 8) containing 10 mM Reduced L-Gluthatione (Sigma-Aldrich, G6529). The sample was dialysed into TBS overnight and the GST-fusion construct was cleaved using Thrombin (Novagen, Merch 69671). dArc capsids formed spontaneously and the cleaved GST protein was removed by washing the sample over a 100 kDa MWCO spin filtration column (GE vivaspin).

### Cryo-EM

5µl of dArc1 or dArc2 capsids at a concentration of ∼1 mg/ml were applied to glow-discharged continuous carbon lacey grids (Lacey Carbon Films on 300 Mesh Copper Grids, Agar scientific), blotted and plunge-frozen in liquid ethane using a FEI Vitrobot Mark IV. Cryo-EM images were acquired on a 300 keV FEI Titan Krios microscope equipped with a Gatan K2-Summit 4Kx4K detector operated in counting mode. A GIF-quantum energy filter (Gatan) was used with a slit width of 20 eV to remove inelastically scattered electrons. Both the dArc1 and dArc2 datasets were collected using SerialEM (Mastronarde, 2003) at a nominal magnification of 105,000 with calibrated pixel sizes of 1.211 Å for dArc1 and 1.388 Å for dArc2. Micrographs were recorded as movies divided into 75 frames. For dArc1 we used a total exposure time of 6.6 s. and an accumulated dose of 35.4 electrons per Å^2^. For dArc2, the total exposure time used was 16.8 s and the accumulated dose 34.9 electrons per Å^2^. Defocus values ranged from −1.0 to −4.0µm. Data collection parameters are summarized in Table S1.

### Image processing

dArc1 and dArc2 image processing and reconstructions are summarized in figures S1 and S2. Dataset parameters are presented in Table S2. Acquired movies were aligned using MotionCor2 (Zheng et al., 2017) with 5 × 5 patches and applying dose-weighting to individual frames. Contrast transfer function (CTF) parameters were estimated using Ctffind4 (Rohou and Grigorieff, 2015). Particles were automatically picked using RELION 3 (Scheres, 2012; Zivanov et al., 2018) and extracted into 512 × 512 pixel boxes. Extracted particles were subjected to several rounds of 2D classification to remove false picks. 2D classes showed views characteristic for icosahedral symmetry, and icosahedral symmetry (I4) was applied for initial model generation and throughout subsequent 3D classification and refinement. A small number of further particles were discarded based on 3D classification. The 3D classification for dArc2 considered only at the CA capsid layer. The dArc2 capsids are less stable at high concentration, and this approach retained more data and produced a higher resolution density after refinement. After refinement we performed per-particle CTF estimation, Bayesian polishing and Ewald sphere correction (Zivanov et al., 2019). The effective resolutions of the cryo-EM density maps were estimated by Fourier shell correlation (FSC = 0.143) (Fig. S1) according to the definition of Rosenthal and Henderson (Rosenthal and Henderson, 2003).

To further improve the resolution, we performed symmetry expansion as implemented in RELION, to calculate the positions and orientations for each of the 2,260,000 asymmetric units for dArc1 and 106,740 for dArc2, centered either at the five-fold or at the two-fold capsomeres. We extracted individual capsomeres using a box sizes of 148 pixels in all cases. For the five-fold capsomeres we removed the redundant five-fold symmetrized capsomeres leaving 453,276 particles for dArc1 and 21,348 particles for dArc2 which were further refined using C5 symmetry. For the dArc1 two-fold capsomeres we performed 3D classification without any applied symmetry and selected only classes displaying clear two-fold symmetric inner density for further refinements (1,685,349 particles). For dArc2 we did not observe any notable differences in classes from the 3D classification, and all particles were used for further refinement. This local refinement approach yielded structures with resolutions of 2.8 Å for the dArc1 capsomeres and 3.7 Å for the dArc2 capsomeres (Fig. S2). For all maps, local resolutions were calculated using ResMap (Kucukelbir et al., 2014) (Fig. S3) and the maps were locally sharpened using LocalDeblur (Ramírez-Aportela et al., 2018).

### Modelling

Model building was performed into the final locally-refined maps. The dArc CA domains were initially modelled based on the NMR structure of the rat Arc CA domain (PDB: 6GSE) (Nielsen et al., 2019) using MODELLER (Webb and Sali, 2017) and the flexible linker between the NTD and CTD as well as the N-terminal tails were *de novo* built using Coot (Emsley and Cowtan, 2004). The Coot models were then refined into the EM density using Phenix real-space refinement (program phenix.real_space_refine) (Adams et al., 2010; Afonine et al., 2018). The dArc1 zinc finger was initially modelled based on the HIV-1 zinc finger (PDB:1A1T) (De Guzman et al., 1998) using MODELLER, rigid-body fitted into the EM density in Chimera (Pettersen et al., 2004) and used as starting point for building additional amino acids in each direction using Coot. We refined the zinc fingers into the EM density using Zen implemented within PDB_REDO (Touw et al., 2016) for ideal tetrahedral coordination of the zinc molecules. Side-chain rotamers and potential clashes at points of contact between the zinc fingers and CA were manually fixed in Coot. Models built into the locally-refined maps were then fitted as rigid bodies into the lower-resolution full capsid maps. Residues making contacts between capsomeres were refined manually. The quality of all models were evaluated using MolProbity, Ramachandran statistics and Clashscore (Williams et al., 2018), both before and after fitting the into the lower resolution full capsid maps (Table S3).

### Alignments

All Alignments were performed using T-COFFEE (Notredame et al., 2000) and figures prepared using ESPript 3.0 (Robert and Gouet, 2014). Input sequences and PDB files are provided in Table S4.

---

**Figure S1.**
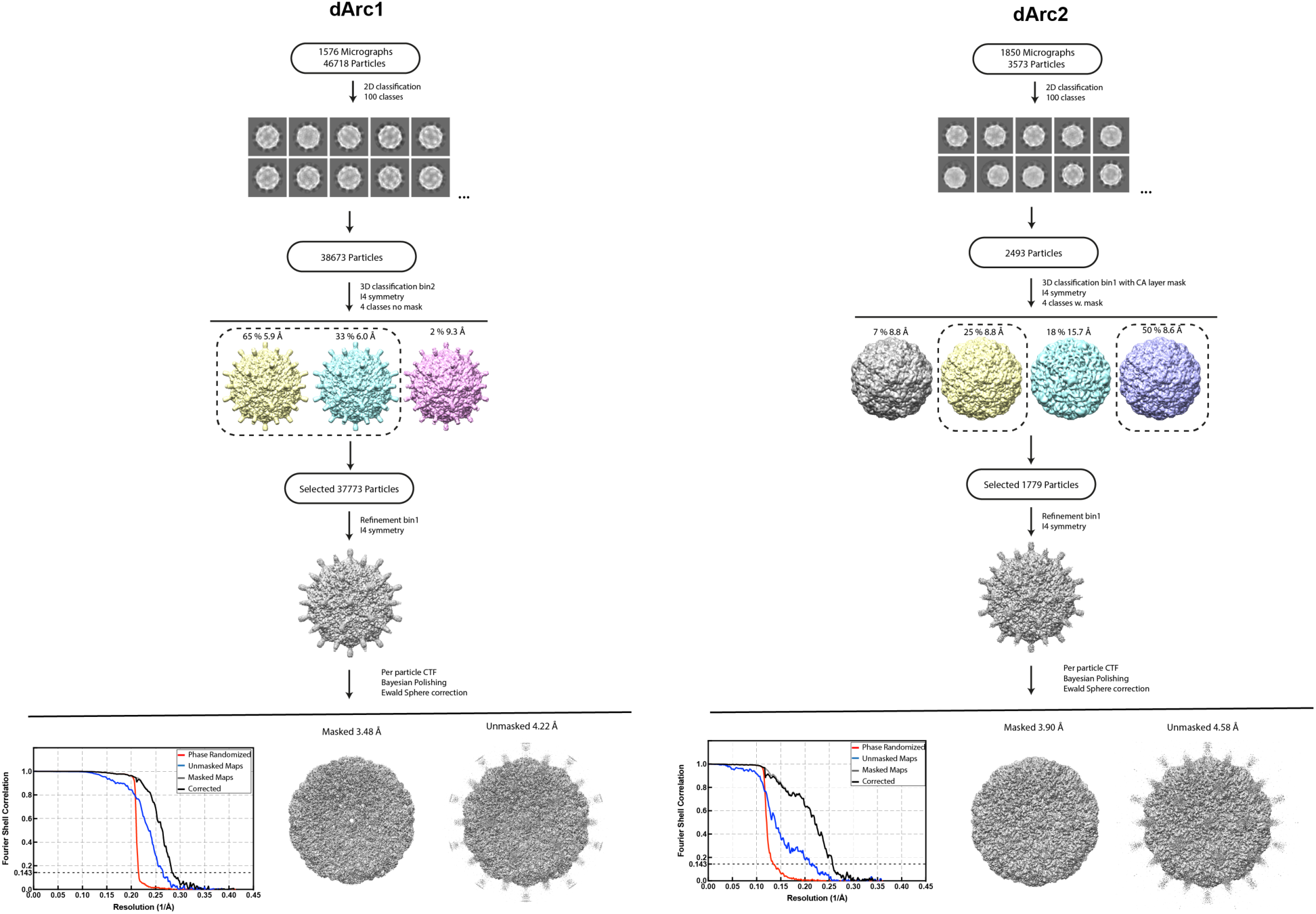
Image processing flowcharts for initial icosahedral dArc1 and dArc2 reconstructions. For details see materials and methods. Resolution of reconstructions are determined by gold-standard Fourier shell correlation (FSC) at the 0.143 criterion.

**Figure S2.**
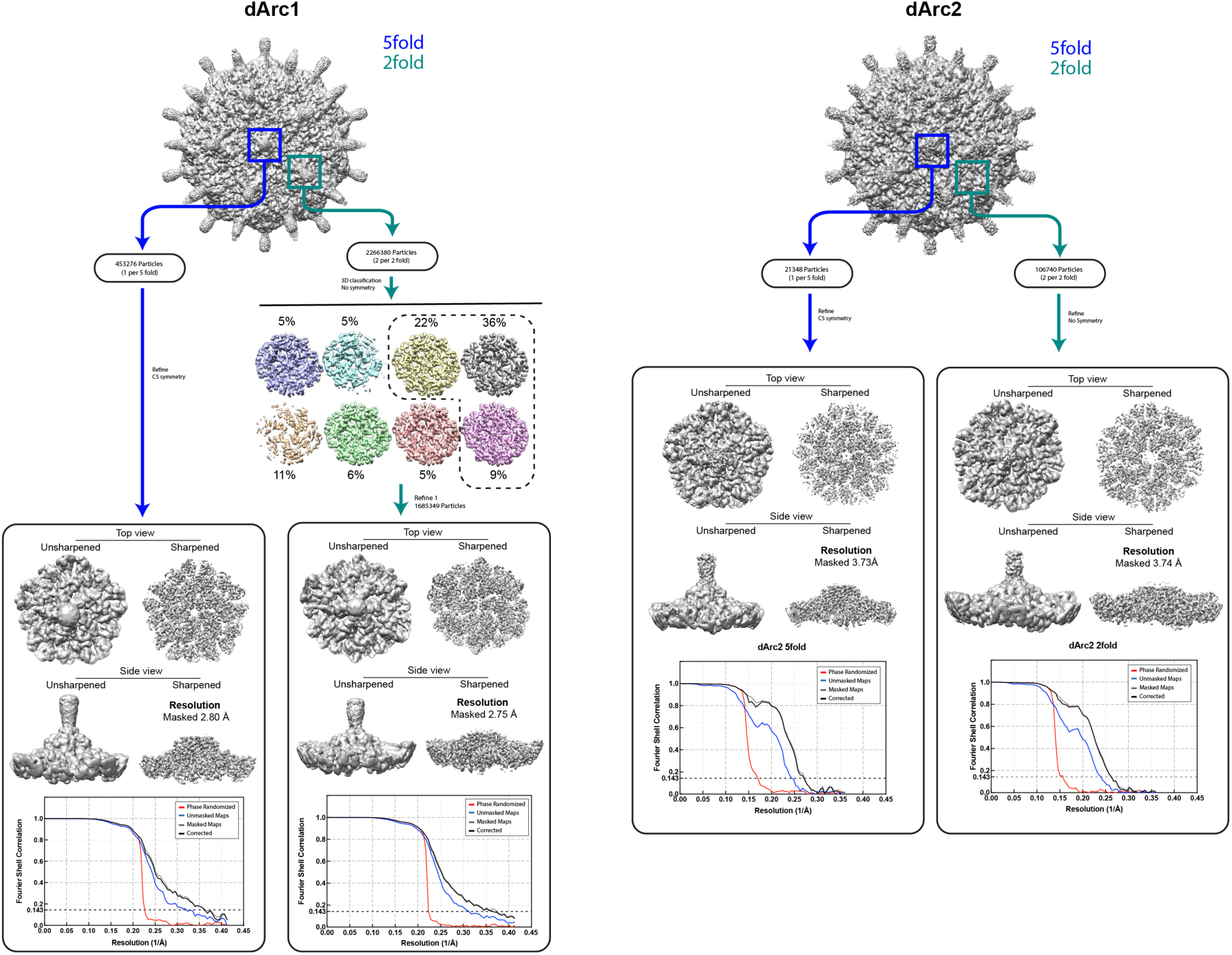
Image processing flowcharts for dArc1 and dArc2 localized reconstructions of five- and two-fold capsomeres. For details see materials and methods. Resolution of reconstructions are determined by gold-standard FSC at the 0.143 criterion.

**Figure S3.**
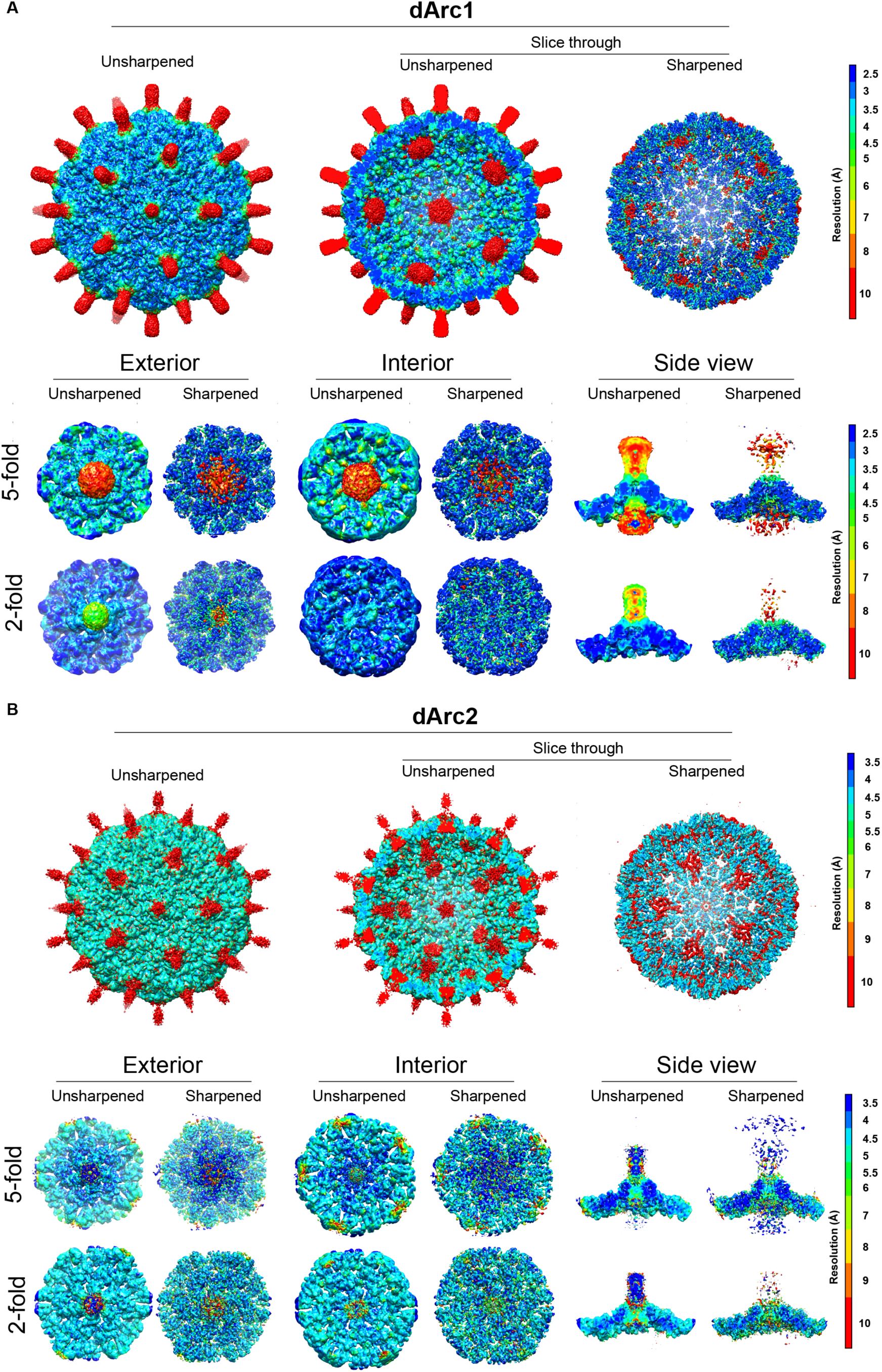
dArc1 and dArc2 density maps colored according to local resolution. Local resolutions were calculated using ResMap(Kucukelbir et al., 2014) and subsequently used for local sharpening of the maps using LocalDeblur(Ramírez-Aportela et al., 2018). **A)** The unsharpened and sharpened maps of dArc1 capsid (top) and locally-refined capsomeres (bottom) colored from high (blue) to low (red) local resolution. **B)** As in A for dArc2. In both cases the CA capsid shells are very well resolved compared to the protruding spikes and the central inner densities. The zinc finger density beneath the dArc1 two-fold capsomeres is well resolved.

**Figure S4:**
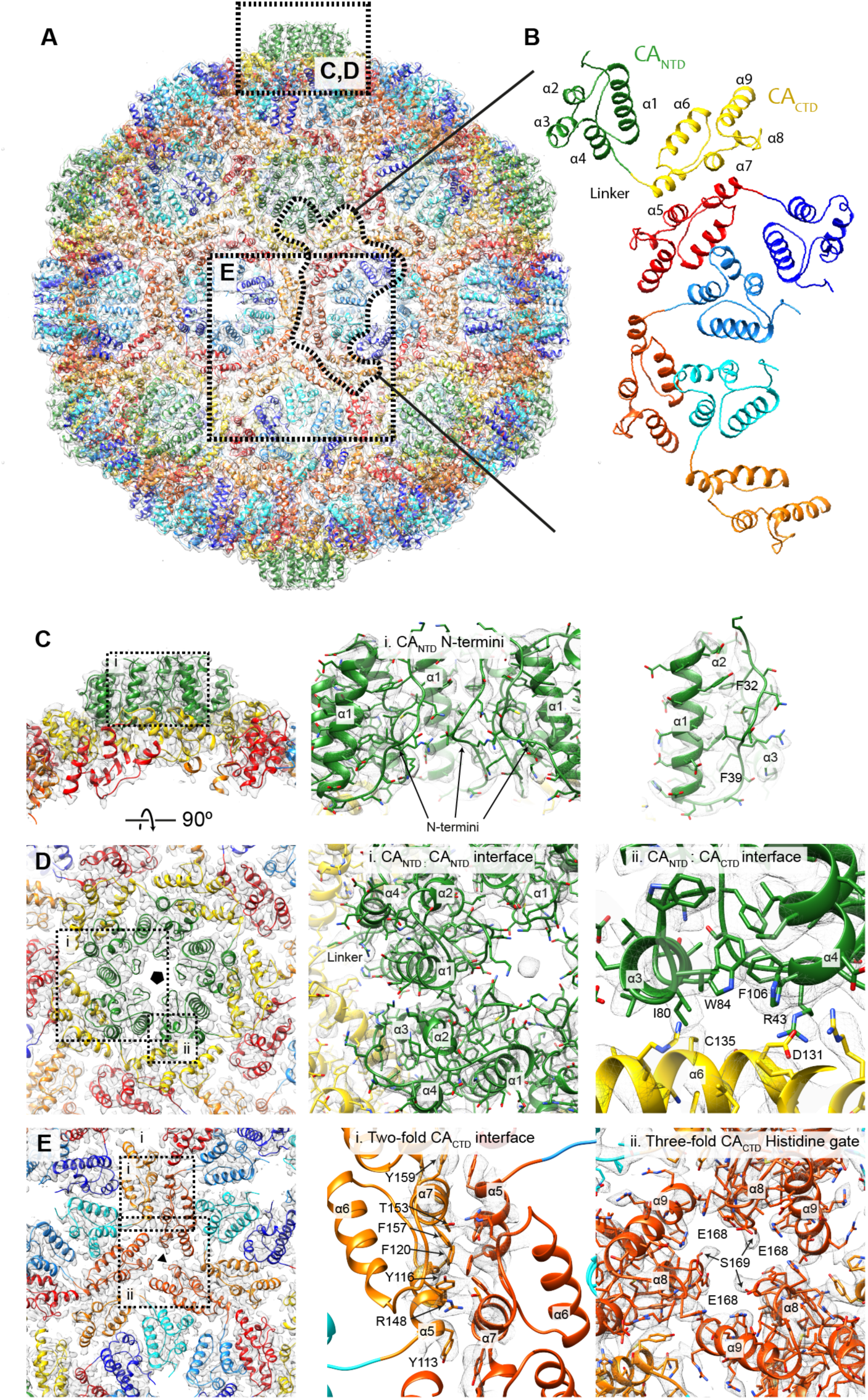
Full capsid atomic model of the dArc2. The views and color scheme are similar to those shown in figure 2. **A)** The 12 five-fold capsomeres are coloured in green (CA_NTD_), and yellow (CA_CTD_). The 30 two-fold capsomeres are coloured in cyan to blue (CA_NTD_) and orange to red (CA_CTD_). **B)** The asymmetric unit containing four CA molecules. 60 asymmetric units including 240 individual CA molecules make up the T=4 capsid. **C)** Close-up of the five-fold capsomere (outlined in A) i) Cut-away showing three out of five N-termini in the centre of the capsomeres. The N-termini extend into and form the capsid spikes. The N-termini are stabilized by docking into an extended hydrophobic groove adjacent to *α*1. **D)** External view of the five-fold capsomere i) The CA_NTD_:CA_NTD_ interaction between *α*1, and *α*2 and *α*3 of the neighbouring CA molecule in the capsomere. Electronegative charged residues on the outside of *α*1 interact with electropositive charges in *α*2 and *α*3. ii) The CA_NTD:_CA_CTD_ interface which involves *α*6 in the CA_CTD_ and *α*3 and *α*4 in the neighbouring CA_NTD_. This interface relies on both hydrophobic and electrostatic interactions. Residues R43, I80, W84, F106, C135 and D131 are depicted in the figure. **E)** External view of the two and three-fold CA_CTD_ interfaces. i) The two-fold CA_CTD_ interface connects two adjacent capsomeres and is dominated by hydrophobic *π* stacking interactions. The interface involves residues from *α*5 and *α*. Residues Y113, Y116, F120, R148, T153, F157 and Y159 are depicted in figure. EM density is shown only for the contact site. All CA_CTD_ interfaces are highly similar (Fig. S7). ii) The three-fold CA_CTD_ axis. At this position, instead of the histidine present in dArc1, E168 and S169 from *α*8 surround the largest gap in the capsid. The corresponding views for dArc1 are shown in Fig. 2.

**Figure S5.**
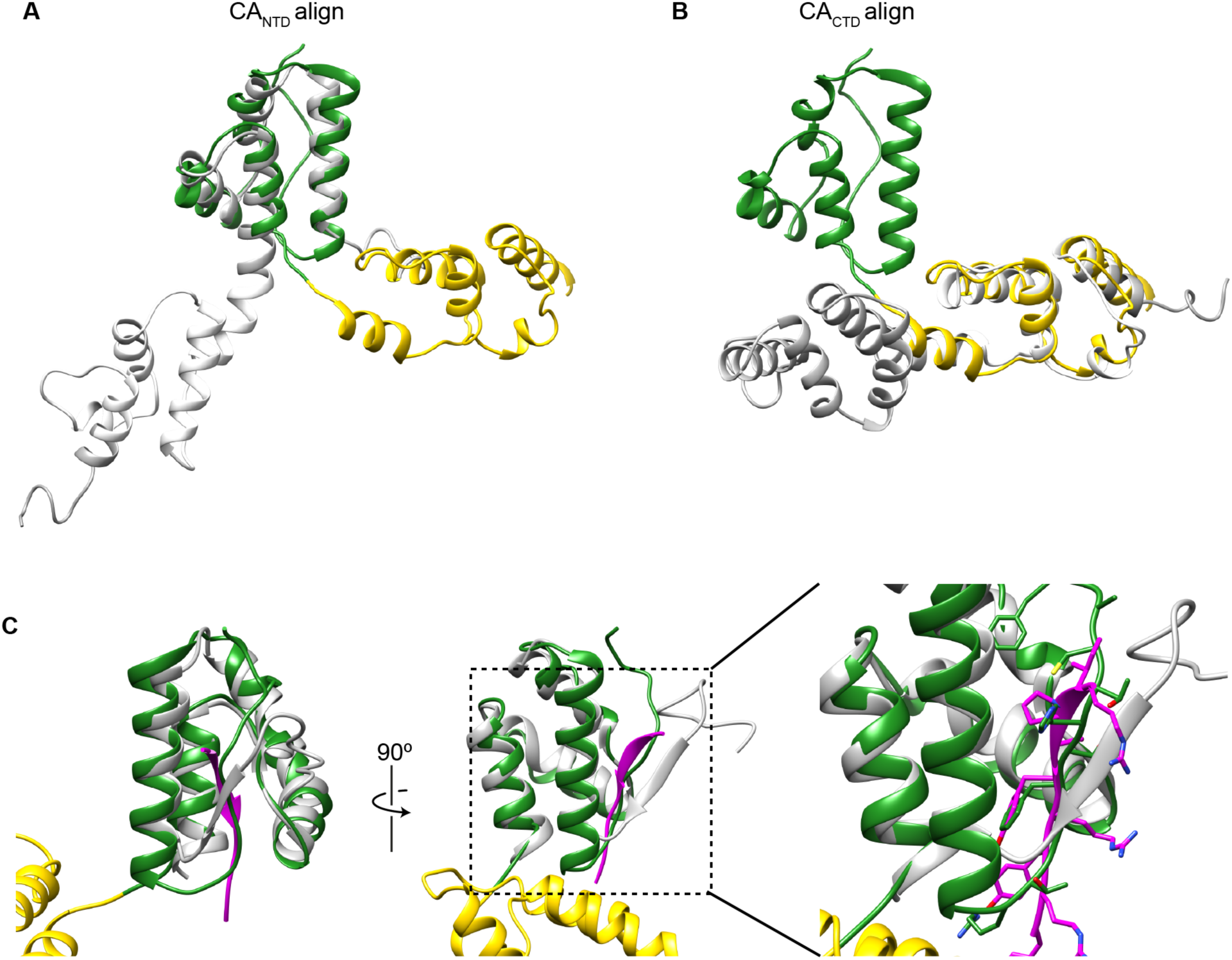
Alignment of the dArc1 and rat Arc structures. **A-B)** The dArc1 CA structure from the five-fold capsomere, compared to the full-length rat Arc (rArc) CA structure (obtained by Nuclear Magnetic Resonance, PDB ID: 6GSE) (Nielsen et al., 2019). The rArc CA_NTD_ and CA_CTD_ are depicted in grey and light grey, respectively. The individual CA_NTD_ and CA_CTD_ folds are completely conserved. The flexible linker connecting CA_NTD_ and CA_CTD_ in the dArc CA capsid structure is more rigid in the monomeric rArc and the interdomain orientation is different. **C)** Alignment between dArc1 CA_NTD_ (green) and the rArc CA_NTD_ (grey) crystal structure (PDB ID: 4X3H) (Zhang et al., 2015) bound to the transmembrane AMPAR regulatory protein *γ*2 (TARP*γ*2) (pink). The binding site for TARP*γ*2, CaMKII and NMDA peptides in rArc, is occupied by the N-terminus in dArc1 and dArc2.

**Figure S6.**
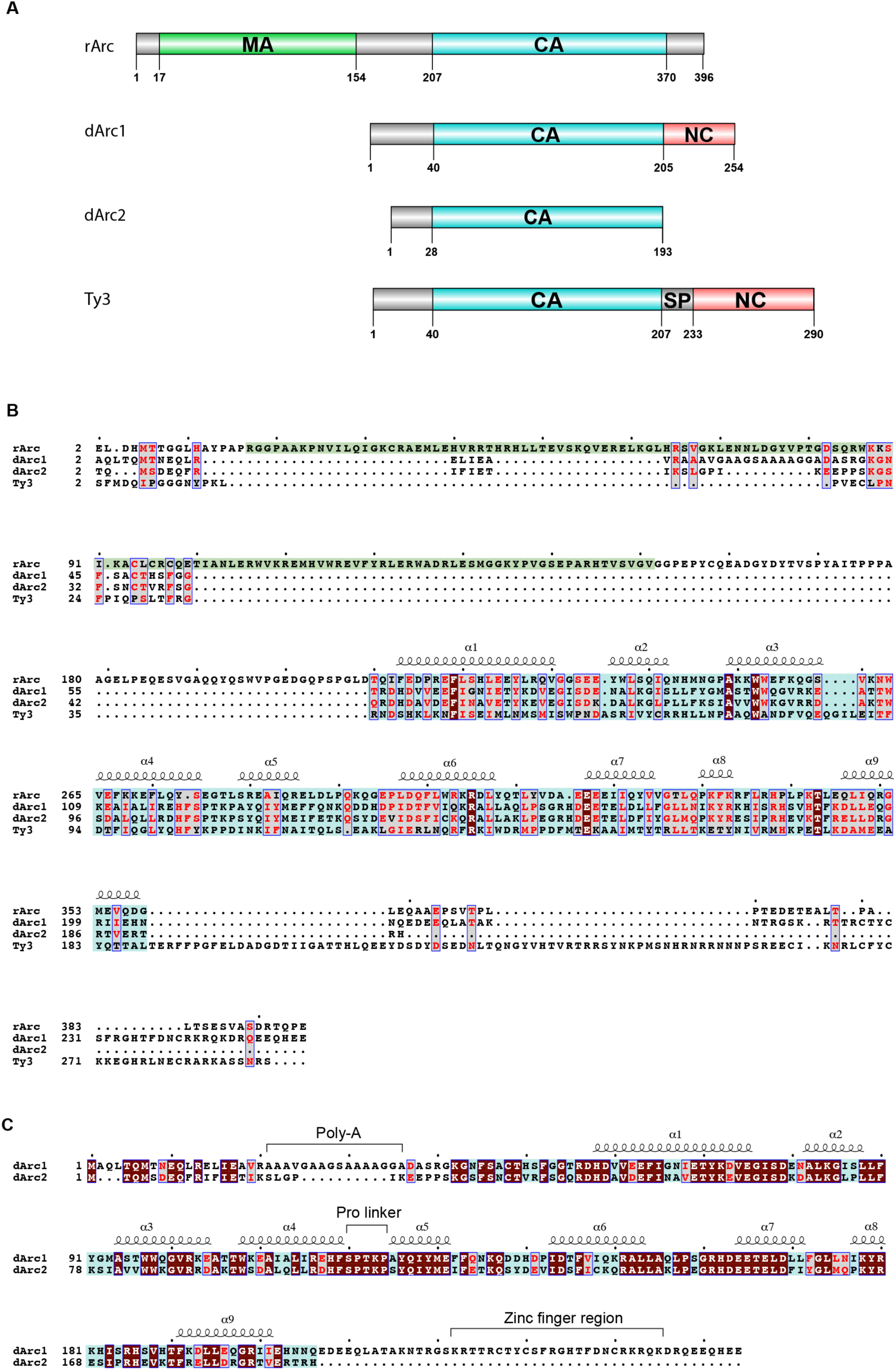
Sequence alignment of dArc1 and dArc2, rArc and Ty3. **A)** Domain overview. Matrix-like domain (MA; Green), capsid domain (CA; blue), nucleocapsid domain (NC; red). Only rArc contains a putative MA domain but lacks the NC domain. dArc2 is shortest of the four and only codes for the CA domain. **B)** The amino acid sequence alignment shows good overall alignment of the CA coding region for all proteins. Conserved residues; Brown, Equivalent residues (T-Coffee equivalence score >0.7); grey. **C)** The amino acid sequence alignment between dArc1 and dArc2. Except for the Poly-A stretch and the NC domain, the sequences are highly conserved. The zinc finger region is marked.

**Figure S7.**
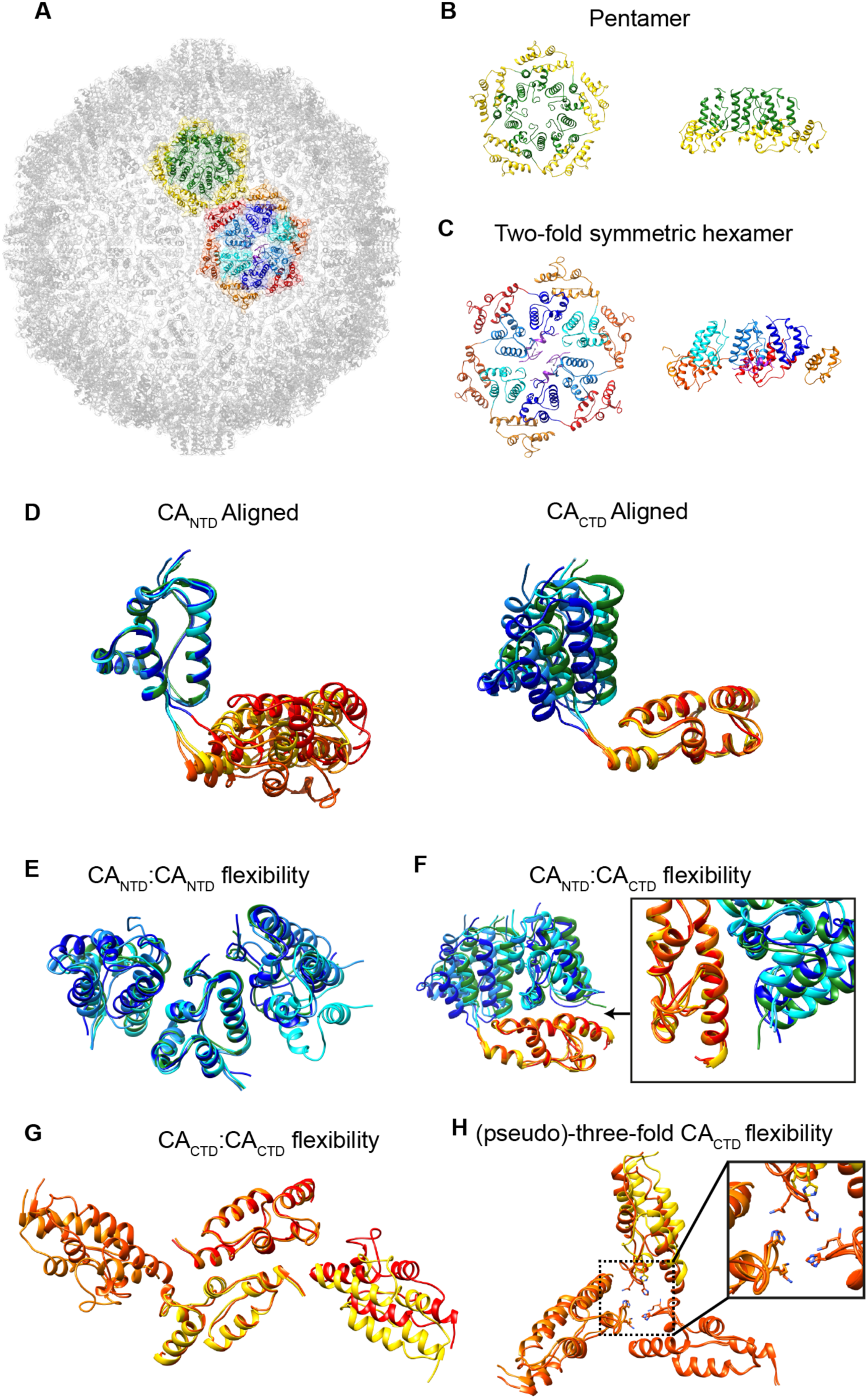
Flexibility of the CA domain within the dArc1 capsid. **A)** Overview of the atomic structure of the dArc1 capsid with one five-fold and one two-fold capsomere highlighted. **B-C**) Top and side views of the five-fold and two-fold capsomeres to indicate the color coding in the following panels. **D)** The four different conformations of CA in the asymmetric unit aligned by either the CA_NTD_ or the CA_CTD_ domain. **E)** Flexibility of the four different CA_NTD_:CA_NTD_ interfaces in the capsomeres, with the central CA_NTD_ aligned. **F)** Flexibility of the four different CA_NTD_:CA_CTD_ interfaces, with the CA_CTD_ aligned. There are only subtle movements of the neighboring CA_NTD_ relative to the CA_CTD._ **G)** The two different conformations of the CA_CTD_:CA_CTD_ interface. The CA_CTD_ domains forming the interfaces between adjacent capsomeres are less variable than the interfaces created between CA_NTD_:CA_CTD_ and CA_NTD_:CA_NTD_ shown above. **H)** Alignment of the three-fold CA_CTD_ and pseudo-three-fold CA_CTD_ axes. The histidines are positioned 6 Å apart in the true three-fold and 6 or 10 Å apart in the pseudo three-fold axes.

**Figure S8:**
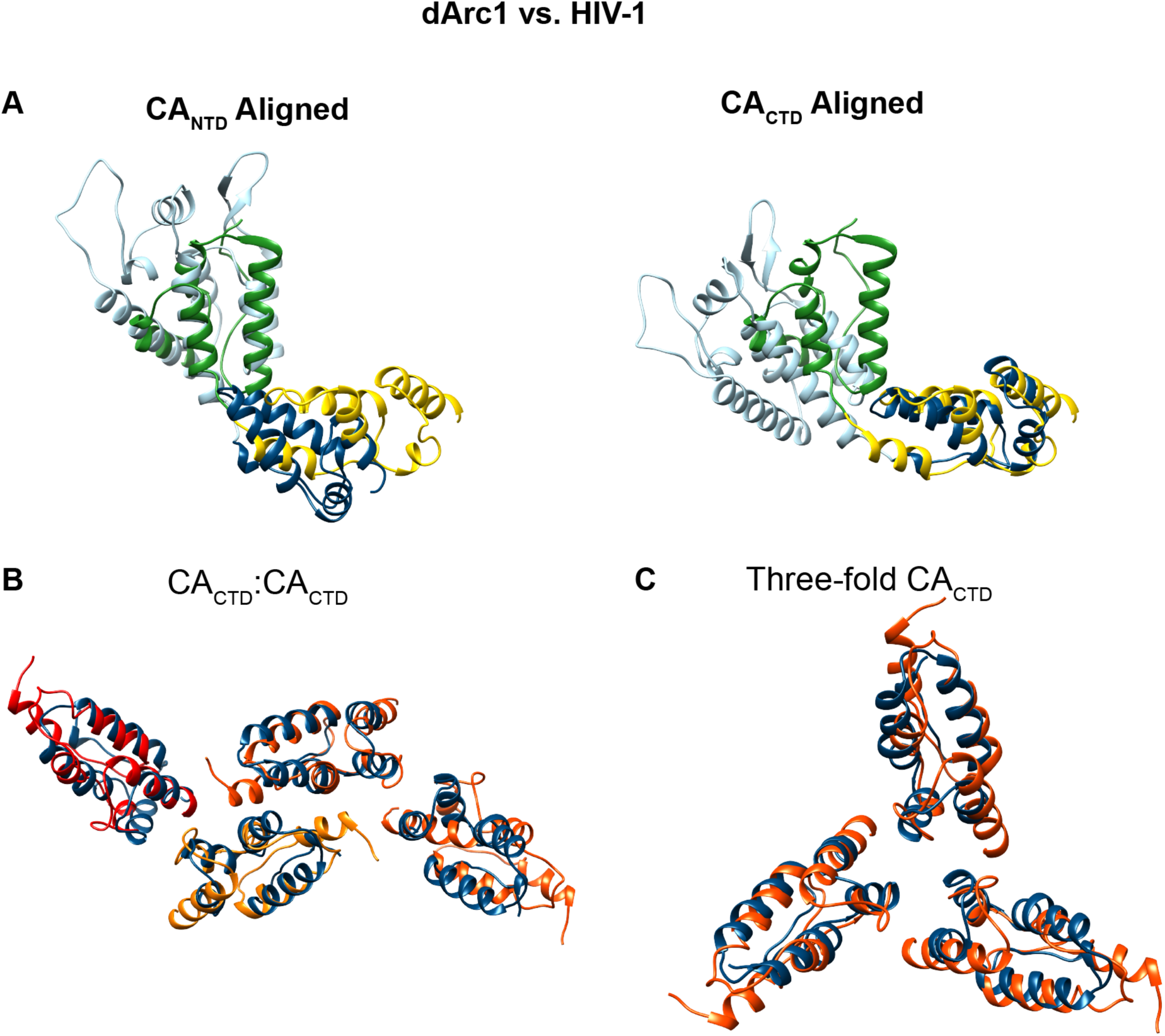
Alignment of the dArc1 and mature HIV-1 structures. **A)** CA from the five-fold capsomere of the dArc1 capsid aligned with CA from the HIV-1 CA pentamer from mature virions (PDB ID: 5MCY) (Mattei et al., 2016) by either CA_NTD_ (HIV-1; light blue) or CA_CTD_ (HIV-1; dark blue). The HIV-1 CA_NTD_ is composed by 7 helical segments and an N-terminal β-Hairpin. *α*1-*α*4 in dArc CA_NTD_ correspond to *α*2-*α*4 and *α*7 in the HIV-1 CA_NTD_. The CA_CTD_ fold is well conserved. **B-C)** The arrangement of CA_CTD_ at the two and three-fold interfaces in the dArc capsid is similar to the arrangement at the corresponding positions in mature HIV-1 (PDB ID: 5MCX) (Mattei et al., 2016). The HIV-1 two-fold CA_CTD_ interface is constituted by a short 3_10_ helical segment and *α*9 which align with dArc *α*5 and *α*7, respectively. Mutations in HIV-1 CA *α*9 are known to disrupt virus assembly and maturation (Joshi et al., 2006; Schwedler et al., 2003).

**Figure S9:**
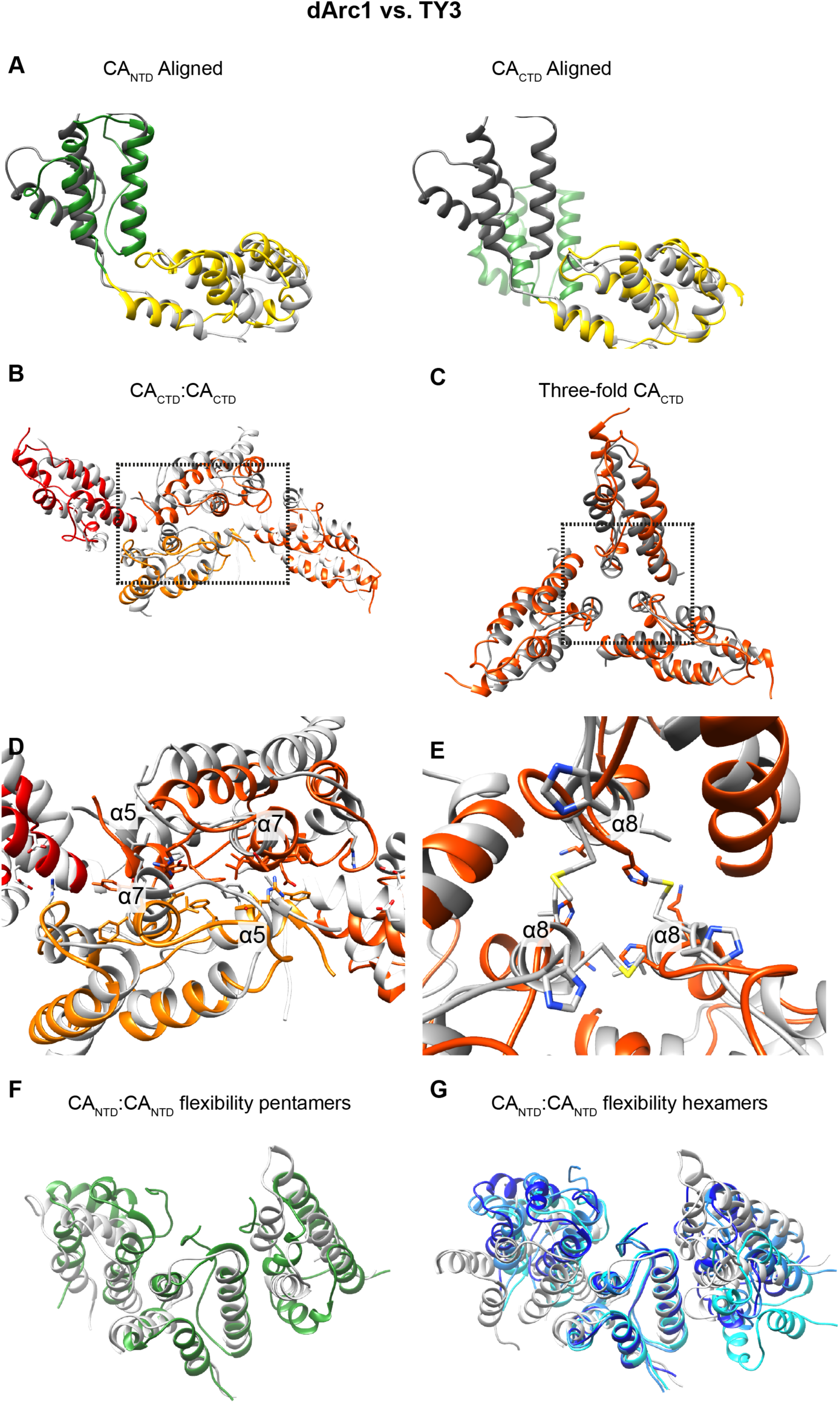
Alignment of the dArc1 and Ty3 structures. **A)** CA from the five-fold capsomere of the dArc1 capsid aligned with CA from the five-fold capsomere of Ty3, either aligned on CA_NTD_ (Ty3; dark grey) or CA_CTD_ (Ty3; light grey). The structures of CA_NTD_ and the CA_CTD_ domains are very similar but differ in the inter-domain orientation. **B-C**) The arrangement of CA_CTD_ at the two and three-fold dArc interfaces is very similar to the arrangement at the corresponding positions in Ty3. **D)** The two-fold CA_CTD_: CA_CTD_ contact surfaces for dArc1 and dArc2 are facilitated by hydrophobic stacking interactions, primarily between *α*5 and *α*7. The Ty3 interface also involves hydrophobic residues in *α*5 and *α*7, but lacks direct *α*5 - *α*5 contacts. **E)** The Ty3 three-fold CA_CTD_ interface also contains a three-fold symmetric histidine cluster (H170) positioned at the outer edge of the pore. **F)** Comparison of dArc1 (green) and the Ty3 (grey) CA_NTD_:CA_NTD_ interfaces from pentameric capsomeres aligned by the central CA_NTD._ **G)** Comparison of dArc1 (shades of blue) and the Ty3 (shades of grey) CA_NTD_:CA_NTD_ interfaces from hexameric capsomeres aligned by the central CA_NTD_.

**Figure S10.**
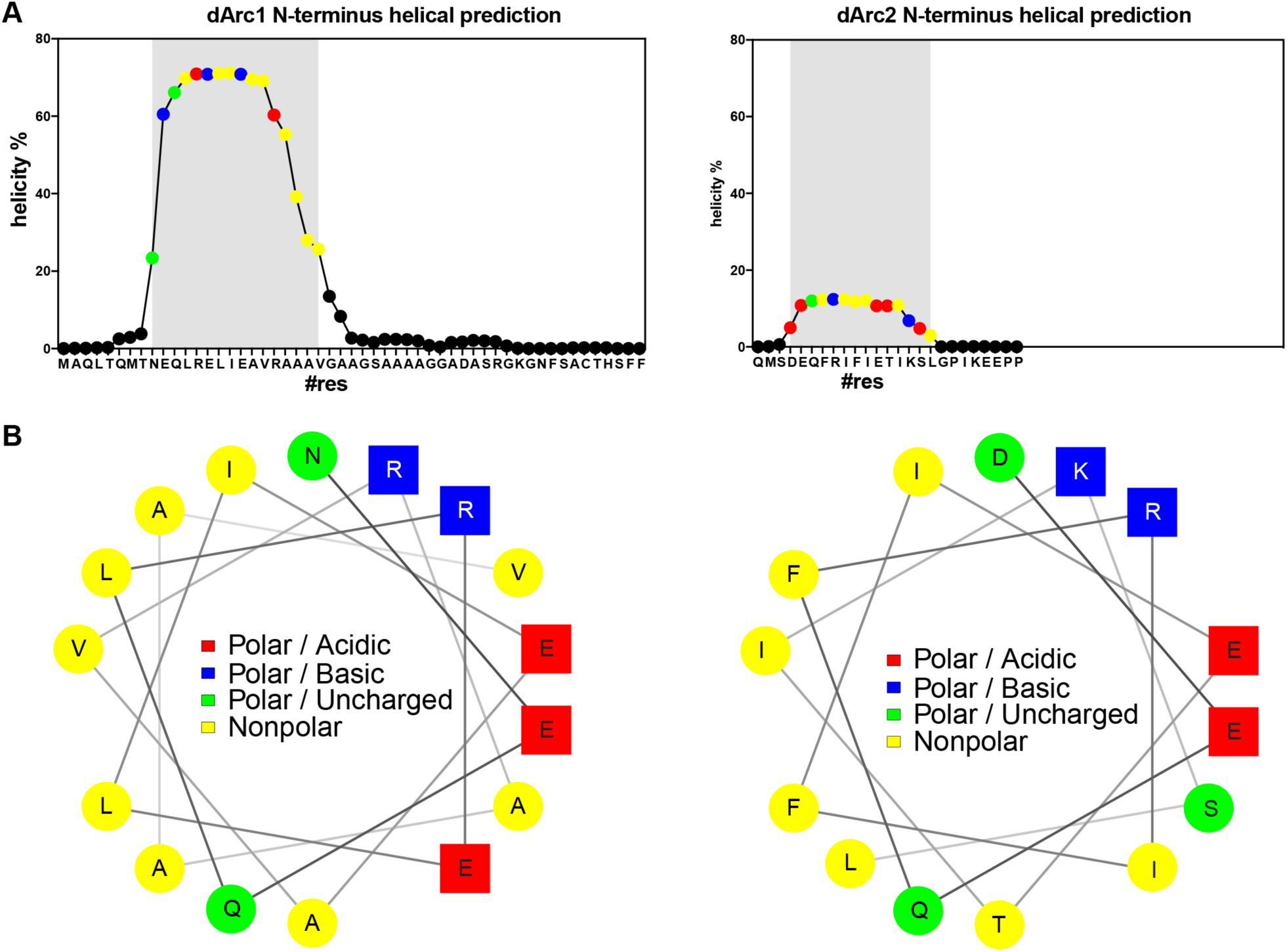
Helical and amphipathic predictions for the dArc N-termini. **A)** Agadir (Muñoz and Serrano, 1994) helical propensity prediction of the 41 residues N-terminal to dArc1 CA and the 28 residues N-terminal to dArc2 CA. **B)** Helical wheel projections showing that the predicted dArc1 and dArc2 helices would be amphipathic.

**Figure S11.**
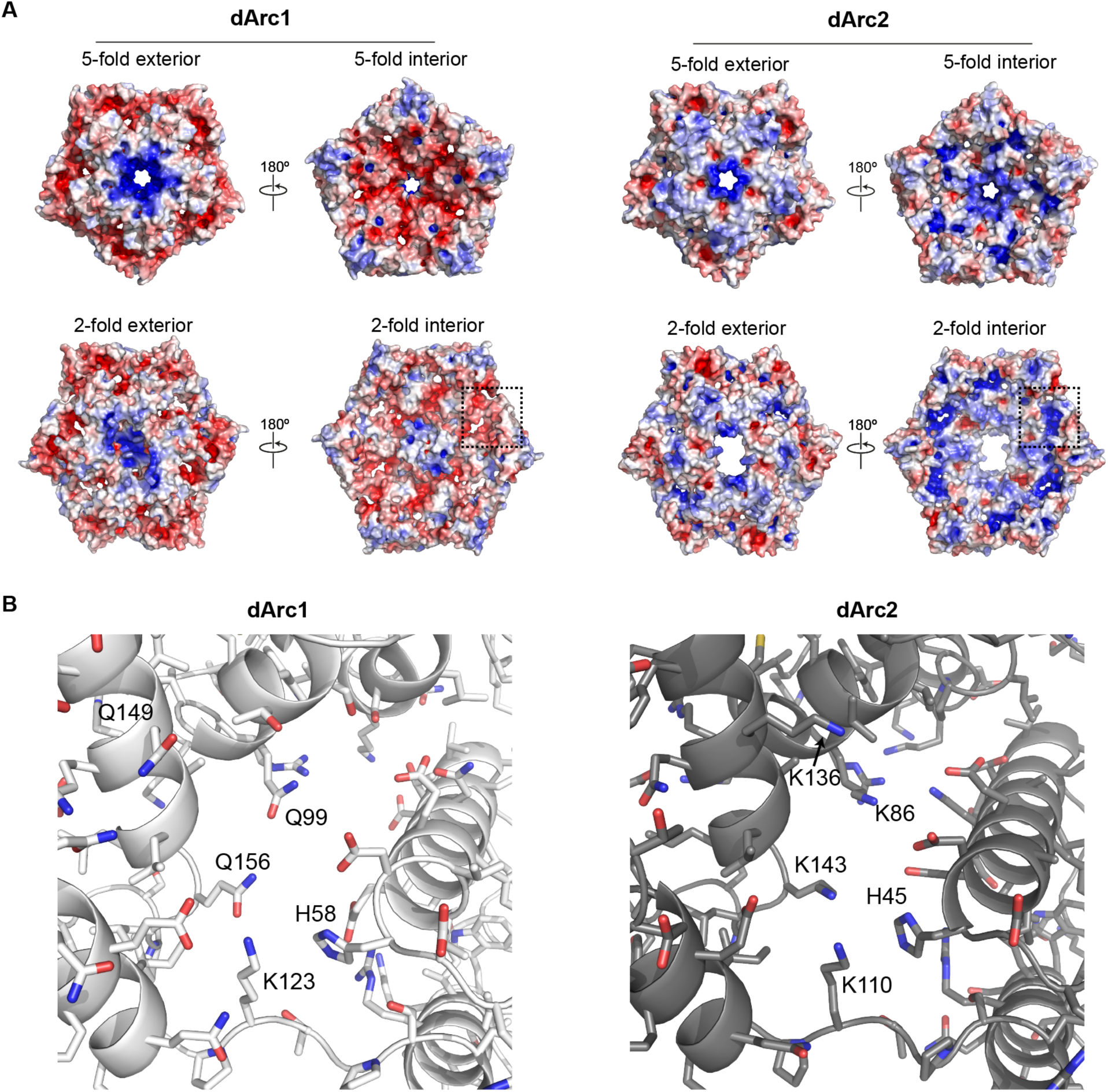
Electrostatic potential of individual capsomeres of dArc1 and dArc2. **A)** As in Fig 3, the surface is coloured with the electrostatic potential from -5 (red) to +5 k_b_T/e (blue). The dArc2 capsomeres are more electropositive than the dArc1 capsomeres. **(B)** Close-ups showing that additional neutral-to-basic residue substitutions facilitate electrostatic charge differences in the capsid interior.

**Table S1:** Data Acquisition

| **Table S1: Data Acquisition** |  |  |
| --- | --- | --- |
|  | **dArc1** | **dArc2** |
| Microscope | FEI Krios | FEI Krios |
| Voltage (KeV) | 300 | 300 |
| Camera | K2 summit (Gatan) | K2 summit (Gatan) |
| Nominal Magnification | 105,000 | 105,000 |
| Defocus range ($\mu\text{m}$) | -1 to -4 | -0.8 to -3 |
| Frames | 75 | 75 |
| Total Exposure (s) | 6.5625 | 16.875 |
| dose (e/px/s) | 7.915 | 3.989 |
| Pixel Size ($\text{\AA}$) | 1.211 | 1.388 |
| dose (e-/$\text{\AA}^2$/s) | 5.396 | 2.071 |
| Dose per frame (e-/$\text{\AA}^2$) | 0.472 | 0.466 |
| Total electron dose (e-/$\text{\AA}^2$) | 35.4 | 34.9 |

**Table S2:** Data Processing and Reconstruction

| **Table S2: Data Processing and Reconstruction** |  |  |  |  |  |  |
| --- | --- | --- | --- | --- | --- | --- |
|  | **dArc1** |  |  | **dArc2** |  |  |
|  | **Capsid** | **5-fold** | **2-fold** | **Capsid** | **5-fold** | **2-fold** |
| Box size (px) | 512 | 148 | 148 | 512 | 148 | 148 |
| Particles | 37,773 | 453,276 | 2,266,380 | 1,779 | 21,348 | 106,740 |
| Symmetry | I4 | C5 | C1 | I4 | C5 | C1 |
| Resolution (Å, FSC = 0.143) | 3.5 | 2.8 | 2.8 | 3.9 | 3.7 | 3.7 |
| Relion B-factor* | -108 | -130 | -116 | -56 | -142 | -165 |

\*Note, all final density maps are sharpened based on local resolution using ResMap (Kucukelbir et al., 2014) and LocalDeblur(Ramírez-Aportela et al., 2018).

**Table S3:** Atomic Models

| **Table S3: Atomic Models** |  |  |  |  |  |  |
| --- | --- | --- | --- | --- | --- | --- |
|  | **dArc1** |  |  | **dArc2** |  |  |
|  | **Asym. unit** | **5-fold** | **2-fold** | **Asym. unit** | **5-fold** | **2-fold** |
| Composition |  |  |  |  |  |  |
| Atoms (non Hydrogen) | 5620 | 6715 | 8554 | 5436 | 6795 | 8154 |
| Ligands | Zn (1) | - | Zn (2) | - | - | - |
| Bonds (RMSD) |  |  |  |  |  |  |
| Length (Å) (# > 4σ) | 0.006 (0) | 0.005 (0) | 0.006 (0) | 0.004 (0) | 0.005 (5) | 0.004 (0) |
| Angles (°) (# > 4σ) | 0.856 (8) | 0.84 (0) | 0.775 (9) | 0.893 (4) | 0.892 (10) | 0.89 (6) |
| MolProbity score | 2.03 | 1.72 | 2.05 | 1.78 | 1.39 | 1.56 |
| Clash score | 10.07 | 4.32 | 10.31 | 6.49 | 1.85 | 3.58 |
| Rotamer outliers (%) | 0.83 | 0.56 | 0.72 | 0.17 | 0 | 0 |
| Cβ outliers (%) | 0 | 0 | 0 | 0 | 0 | 0 |
| Ramachandran plot (%) |  |  |  |  |  |  |
| Favored | 91.57 | 91.36 | 91.25 | 93.67 | 93.21 | 93.83 |
| Allowed | 7.99 | 8.64 | 8.27 | 6.33 | 6.79 | 6.17 |
| Outliers | 0.44 | 0 | 0.49 | 0 | 0 | 0 |

**Table S4:** Input sequences and PDB files

| **Protein** | **UniProtKB** | **PDB ID** |
| --- | --- | --- |
| Ty3 | Q7LHG5 | 6R24 |
| rArc | Q63053 | 6GSE, 4X3H |
| HIV-1 p24 gag | B6DRA0 | 5MCX, 5MCY |
| HIV-1 Nucleocapsid protein | Q75677 | 1A1T |
| dArc1 | Q7K1U0 | this study |
| dArc2 | Q7JV70 | this study |

